# Nanoconfinement of Microvilli Alters Gene Expression and Boosts T cell Activation

**DOI:** 10.1101/2021.04.19.440349

**Authors:** Morteza Aramesh, Diana Stoycheva, Ioana Sandu, Stephan J. Ihle, Tamara Zünd, Jau-Ye Shiu, Csaba Forró, Mohammad Asghari, Margherita Bernero, Sebastian Lickert, Annette Oxenius, Viola Vogel, Enrico Klotzsch

## Abstract

T cells sense and respond to their local environment at the nanoscale by forming small actin-rich protrusions, called microvilli, which play critical roles in signaling and antigen recognition, particularly at the interface with the antigen presenting cells. However, the mechanisms by which microvilli contribute to cell signaling and activation is largely unknown. Here, we present a tunable engineered system that promotes microvilli formation and T cell signaling via physical stimuli. We discovered that nanoporous surfaces favored microvilli formation, and markedly altered gene expression in T cells and promoted their activation. Mechanistically, confinement of microvilli inside of nanopores leads to size-dependent sorting of membrane-anchored proteins, specifically segregating CD45 phosphatases and T cell receptors (TCR) from the tip of the protrusions when microvilli are confined in 200 nm pores, but not in 400 nm pores. Consequently, formation of TCR nanoclustered hotspots within 200 nm pores, allows sustained and augmented signaling that prompts T cell activation even in the absence of TCR agonists. The synergistic combination of mechanical and biochemical signals on porous surfaces presents a straightforward strategy to investigate the role of microvilli in T cell signaling as well as to boost T cell activation and expansion for application in the growing field of adoptive immunotherapy.

## Introduction

T cell microvilli are formed during the formation of immunological synapse, through which T cells strive to gain maximal interactions with the antigen-presenting cell.^1–3^ Besides their exploratory role in maximizing surface contact to detect cognate antigens,^4^ microvilli can regulate T cell signaling by forming areas with high membrane curvature.^5^ Formation of high membrane curvature is considered to be sensed by triggering biological processes, such as the assembly of signaling microclusters. Formation of high membrane curvatures (200-1000 nm in diameter) supports antigen recognition in T cells, through amplified and sustained signaling at the tip of the protrusions.^1,5^ Recent studies suggest that sporadic nonspecific T cell receptor (TCR) phosphorylation can occur at the tip of microvilli, where CD45 is sterically excluded through the formation of “tight contacts”.^6,7^ However, the underlying mechanism and the biological processes triggered by microvilli formation in the process of T cell activation remain largely unknown.

Understanding signaling processes induced by microvilli formation is important both from a physiological and technological point of view. This becomes obvious in Whiskott-Aldrich Syndrome and Leukemia, where disrupted formation of microvilli leads to compromised function of various immune cells.^8^ Importantly, microvilli formation can be regulated by common anticancer drugs such as cytokines, chemokines or cytoskeletal inhibitors, potentially impairing T cells in being able to efficiently recognize their cognate antigens, thus limiting the efficiency of the therapy.^9,10^ Beside chemical signals, the physical environment of the cell can also regulate microvilli formation, particularly at the nanoscale.^11^ The consequences of T cell microvilli regulation by nanomaterial implants is largely unexplored, which is particularly significant considering the rise of nanotechnology-based immunotherapy approaches and a large pipeline of associated treatments in clinical trials.^12,13^ From an engineering perspective, one may exploit the possibility of engineering T cell functions by regulating microvilli formation *ex vivo*, which can have an impact in the growing field of adoptive T cell therapy.

Despite the importance of microvilli as a structural unit of the cell and their role in T cell signaling, there is a limited amount of relevant studies that investigated microvilli function in T cell activation.^4,5,8^ This is mainly due to the fact that microvilli are nanoscale projections from the cell membrane displaying high dynamics of formation and retraction, therefore difficult to investigate by standard optical microscopy.^4,14,15^ While electron microscopy enables their visualization, their dynamic behavior still remains largely unresolved.^1^ The fast and transient nature of microvilli protrusions, makes it challenging to detect downstream signaling events. Using lattice light-sheet microscopy and real-time tracing of dynamic microvilli on 2D surfaces, the life-time of the microvilli protrusions is estimated to be roughly 1 min, corresponding to the half-life of T cell–APC contact duration *in vivo*.^15^

To investigate how microvilli interact with nanoscale surface features, we studied the natural behavior of T cells in exploring their environment when in contact with engineered nanoporous materials. Surprisingly, microvilli formation was triggered by presenting a physical cue. The dimensions of the T cell projections are defined by the pore size, if it matches the dimensions of membrane protrusions used by T cells to explore antigen presenting cells. Particularly, confinement of the microvilli into pores with 200 nm in diameter, markedly alters cell signaling cascades and subsequent gene expression, even enabling physical activation of T cells without TCR engagement. We explain this phenomenon by nanoconfinement enforced spatiotemporal segregation of the CD45 phosphatase from the TCR. Physically separated from the inhibitory function of CD45, TCR downstream signaling is enhanced even in the absence of specific TCR agonistic stimuli. Scientifically, this discovery that size-matched nanopores promote microvilli formation provides a platform to study microvilli formation processes and their role in T cell activation. Technologically, this opens unforeseen opportunities to boost T cell activation and expansion *ex vivo* by inducing microvilli formation with specific dimensions, which has potential applications in cell-based immunotherapies.

## Results and discussion

### T cell microvilli induction and stabilization by nanoporous substrates

While microstructured surfaces were shown to induce formation of membrane protrusions in T cells,^16–18^ T cells reside in nanostructured environments, where they explore the nanoscale features of antigen presenting cells. To match with the membrane protrusions used by T cells, we studied the effect of nanotopographical constraints on microvilli formation in non-activated T cells. Anodic aluminum oxide (AAO) was used as substrate to create nanotopographical constraints for T cell microvilli induction. AAO is produced by anodization of aluminum films, which results in formation of quasi-ordered arrays of nanopores with a narrow distribution of pore size and interpore distance (**Figure S1**). Non-porous films, as control samples, were prepared by atomic layer deposition of aluminum oxide on flat glass coverslips.

The attachment of the Pan T cells to the nanoporous surface (without agonistic TCR stimuli) was first visualized by confocal microscopy (**Figure 1**), showing how the cells extend their actin-rich microvilli into the nanopores (**Figure 1a**, also see the z-stack **Movie S1**). The actin-rich protrusions appeared as dotted structures under the microscope (schematically shown in **Figure 1c**). By obtaining a 3D profile of the cells using actin staining, the length and depth of the protrusions were estimated. A more detailed picture of the actin cytoskeleton was obtained by super-resolution microscopy (Structured Illumination Microscopy, SIM), revealing the infiltrated protrusions were mostly concentrated around the periphery of the cell (**Figure 1b**).

**Figure 1.**
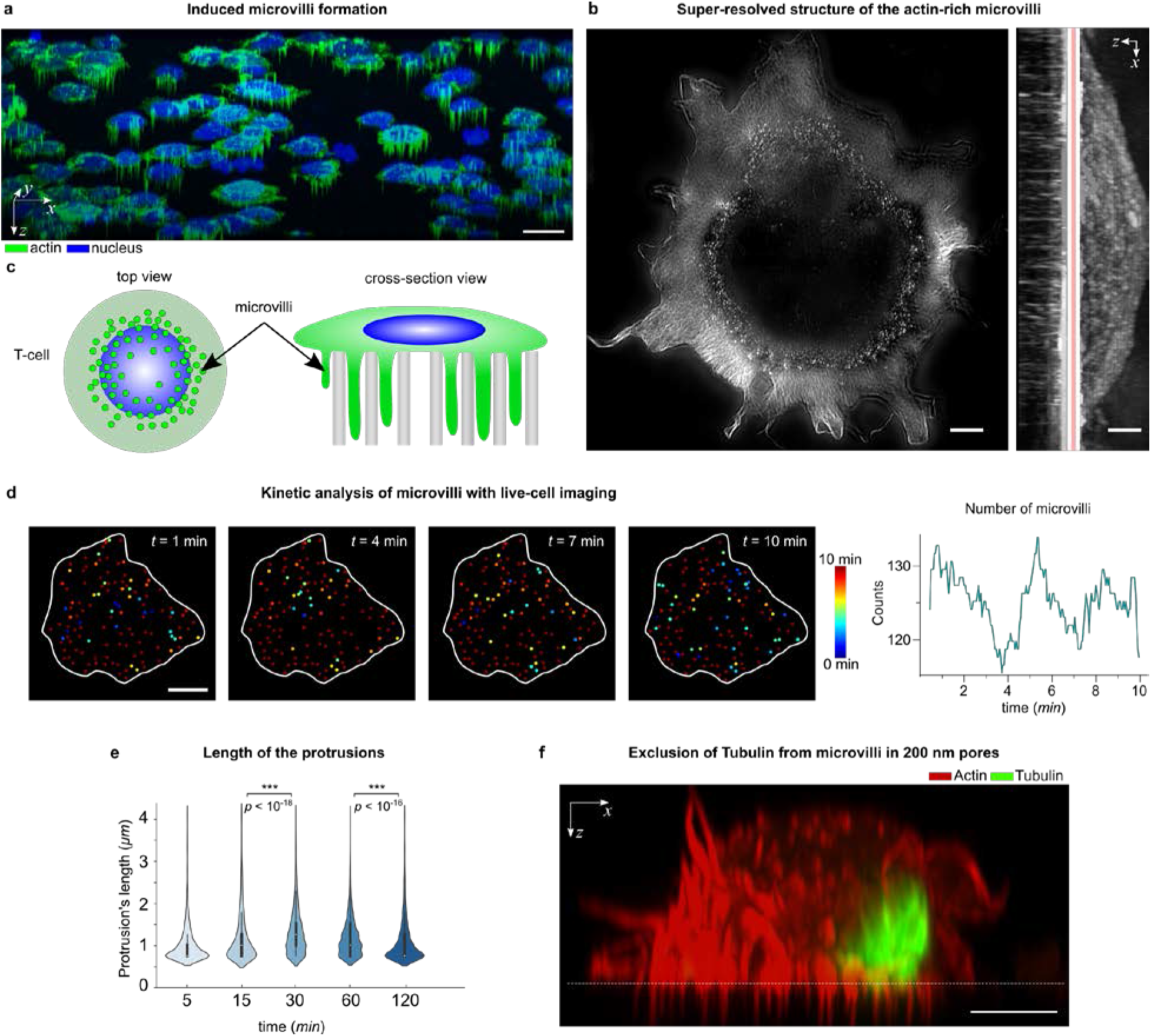
Microvilli protrusions in T cells on porous surfaces. **a)** shows the confocal fluorescence microscopy images of Jurkat T cells on a porous AAO with 200 nm pore size (3D isometric projection), scale bar 10 μm. **b)** shows a super-resolution image of actin cytoskeleton in a T cell using structured illumination microscopy. The protrusions are seen as a dotted structure in the top-view intensity projection and mostly concentrated around the cells’ periphery, scale bar 1 μm. **c)** shows a schematic presentation of the actin-rich protrusions in T cells (not to scale). **d)** (left) shows snapshots of the actin protrusions in a live cell, obtained by fluorescent microscopy. The color corresponds to the lifetime of each protrusion, scale bar 5 μm. (right) shows the number of protrusions in each frame, for the duration of 10 min.**e)** The protrusions’ length was measured on fixed cells at the indicated time points. Violin plots show the distribution of the measured lengths (n = 20 cells). The p-values were determined by two-sided Mann-Whitney tests using R. **f)** A cross--section view of a Jurkat T cell on a porous AAO with 200 nm pore size, captured by Airyscan confocal microscopy. The cells were stained with phalloidin (actin cytoskeleton) and anti-tubulin (microtubules), indicating that the microtubules were mostly excluded from entering the 200 nm pores, scale bar 5 μm.

The microvilli formation occurred very fast and within the first minute after contact with the porous surface (without agonistic TCR stimuli), as seen in **Movie S2**, which was recorded by live cell imaging of Jurkat T cells transfected with Lifeact-GFP, for fluorescently tagging the actin cytoskeleton. The appearance of the bright dotted structure in 2D images is an indication of microvilli formation inside the pores, therefore allowing for the estimation of formation/retraction kinetics. As an example, we collected snapshots (every 10 sec) of the microvilli just below the surface, showing that the formation or retraction of the microvilli is a highly dynamic process with an estimated average rate of ~14 microvilli being formed (and ~13 being retracted) per minute (**Figure 1d** and **Movie S3**). The highly dynamic infiltration of microvilli into the pores enables the cells to maintain their mobility on the surface and to retract from the surface. Retraction of microvilli from the pores was captured by live cell imaging, where we observed *e.g*. detachment of some protrusions at the trailing edge of a migrating cell (**Movie S4**), or gradual yet complete detachment of protrusions upon the cell’s retraction from the surface (**Movie S5**). Moreover, cells can spread their lamellipodia at the interface, and initiate actin treadmilling (**Movie S6**).

By looking into the kinetics of the microvilli’s length over time (*t* = 5, 15, 30, 60 and 120 min), it was observed that the average protrusion length increased with time until *t* = 30 min, reaching to an average length of *L_ave_*~ 1.3 μm, and then decreased after 120 min (*L_ave_*~ 0.9 μm), suggesting that the cells can loosen their contacts to retract from the surface (**Figure 1e**). Notably, protrusions within 200 nm pores are both longer and more stable than those observed on flat surfaces,^15,19^ therefore nanotopography encourages and stabilizes formation of microvilli.

The advantage of using nanoengineered surfaces is that the length, thickness and number of the microvilli can be controlled by tuning the dimensions of the nanotopographical constraints, such as pore depth, pore size and interpore distance. For example, the pore size (and spacing) in AAO films can be finely tuned by anodizing conditions ranging from 20 to 400 nm,^20^ or even smaller via ion-irradiation ^21^, whereas the pore depth can be tuned by anodization time, or by filling the pores with other polymeric materials (**Figure S3**). Surface functionalization can also play a role in microvilli formation, *e.g*. hydrophobic pores were created via silane modification, inhibiting penetration of protrusions into the porous membrane (**Figure S3**).

In order to establish the link between nanotopographical constraints and microvilli dimensions, Jurkat T cells were cultured on AAO membranes with different pore sizes, *i.e*. 20, 100, 200 and 400 nm, as well as on PDMS membranes with 1 μm pores. Notably, T cells were not able to form protrusions on smaller pores, *i.e*. 20 and 100 nm pores, most probably due to the fact that the pores could not physically accommodate microvilli and their protrusive actin-based machinery (**Figure S4**). On the other hand, very large pores – *i.e*. 1 μm pores – are not sufficient to induce microvilli formation. This is particularly evident from the microscope images, where a micron-size pore is only partially infiltrated by a cell protrusion (**Figure S4**), suggesting nanoscale topographical constraints are required to stabilize microvilli. 200 and 400 nm pores were particularly potent to induce microvilli formation. On the mechanistic side, nanopores create a physical barrier – either by size of the pore or by the high membrane curvature at the pore edge – that may exclude some cellular components from entering the pores. Notably, microtubules could not enter the 200 nm pores (**Figure 2e**), whereas they were found in 400 nm pores (**Figure S4**). Microtubules are one of the main tracks for organelle positioning and protein transport.^22,23^

**Figure 2.**
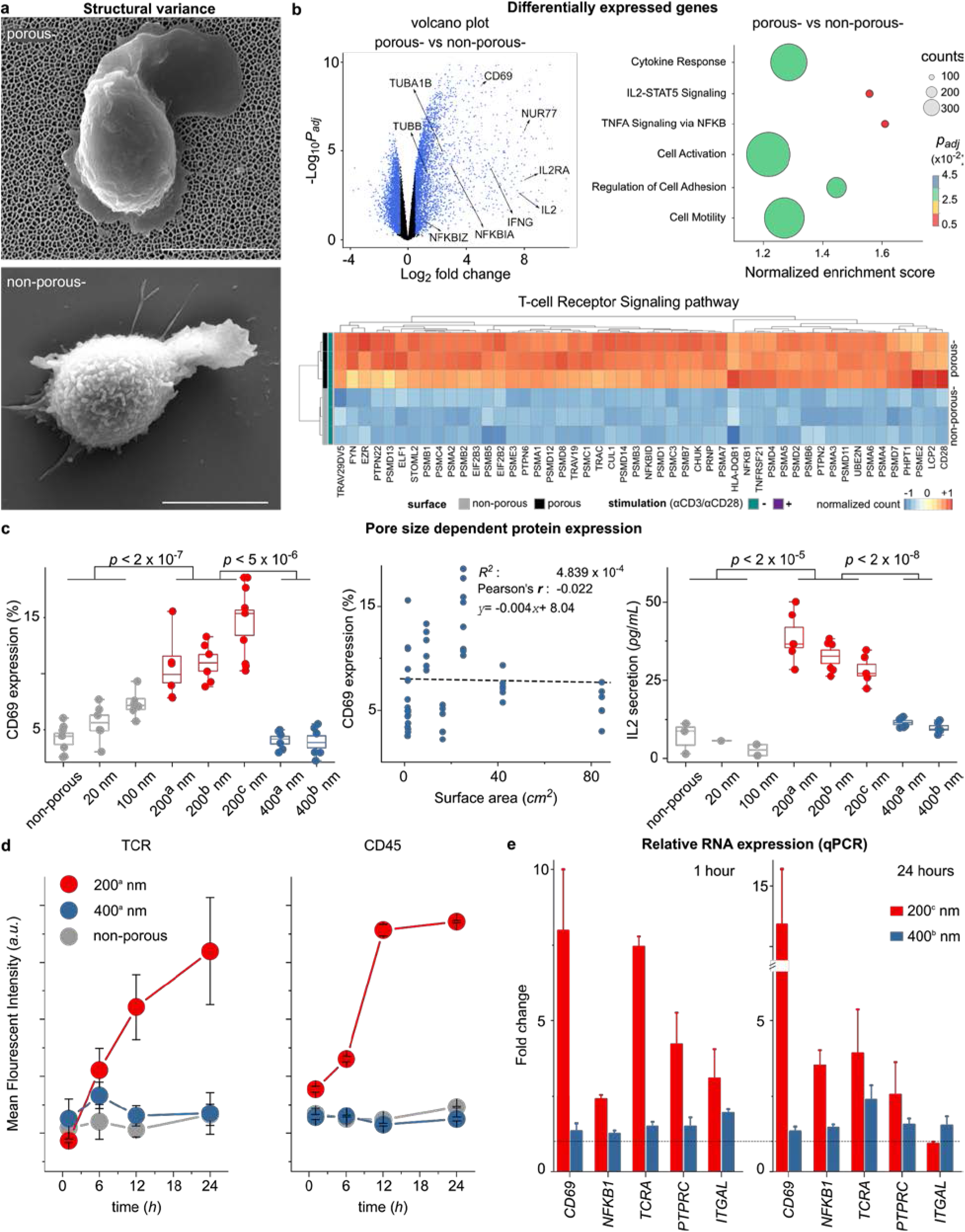
Altered T cell gene expression in 200 nm porous substrates. **a**) shows SEM images of non-stimulated primary human T cell on top of the porous (-) and non-porous (-) surfaces (without αCD3/CD28 coatings), scale bar 4 μm. **b**) Nanoconfinement-induced alteration in the gene program of the primary human T cells after for 4 h of seeding on the surfaces. Volcano plot shows differences in RNA expression of non-stimulated T cells on porous (-) versus non-porous (-) surfaces. Blue color indicates significance with adjusted *P* value < 0.05, and log2(fold changes) > 0.5. Dot plot of gene set enrichment analysis shows differences in differentially enriched pathways of non-stimulated T cells on porous (-) versus non-porous (-) surfaces. The size of the circle corresponds to the gene counts (from the reference pathway), the color corresponds to the adjusted *P* value. (bottom) Heatmap of the top 50 significantly upregulated genes (adjusted *P* values < 0.05) in the TCR signaling pathway, where genes in porous (-) cells were compared to non-porous (-) cells (n = 3 replicates, human primary T cells). The complete set of the significantly differentially expressed genes in TCR signaling pathway can be found in the **supplementary information**. **c)** Jurkat T cells were seeded on porous and non-porous surfaces (without activating antibodies αCD3/CD28) with the indicated pore size (see Table 1). CD69 and IL-2 were measured 24 hours after activation by flow cytometry and Elisa, respectively. Box diagrams show data pooled from two or three independent experiments performed in triplicates. The scatter plot represents the pulled data from all experiments, plotted as surface area versus CD69 expression. The dashed line is the best linear fit to the data, which shows correlation between CD69 expression and increased surface area (Pearson’s r = −0.022). **d)** The scatter plot shows the kinetics of the mean fluorescent intensity (MFI) of TCR and CD45 of primary human T cells cultured on the indicated surfaces at different time points, measured by flow cytometry. Data represent one of two independent experiments performed in triplicates. Error bars are mean ± s.d. **e)** Relative RNA expression of selected genes measured by real-time polymerase chain reaction (RT-qPCR). Primary human T cells were cultured on the indicated surfaces for 1 hour and 24 hours. Fold expression is measured against the control sample (non-porous) and normalized by the reference gene RNA18S. The dashed line indicates fold expression = 1, the expression of the genes in the control sample. The p-values were determined by two-sided Mann-Whitney tests in R.

Similar to the signaling effect of receptor segregation imposed by 2D micropatterns ^24^, we might expect that the exclusion of cellular components from microvilli via nanoscale confinement in pores may have significant biological consequences. For example, vesicles that are normally transported via microtubules to the tip of the protrusions, may not be transported to the confined regions, potentially altering cell signaling. Moreover, size exclusion of membrane proteins can occur, which are very critical for T cell signaling (discussed later).

### Nanoscale microvilli formation alters gene expression and T cell signaling

In order to evaluate whether microvilli nanoconfinement has an effect on signaling of primary human T cells, early gene expression analysis was conducted 4 h after T cell seeding. Naive T cells were incubated on non-porous and porous membranes (200 nm), without the presence of TCR activating antibodies (αCD3/CD28) on the surface, referred to as non-porous (-) and porous (-), respectively (**Figure 2a**). The expression of ~6300 genes distinguished T cells being exposed to porous (-) or non-porous (-) conditions, with ~3300 genes having higher expression in cells being exposed to porous (-) versus non-porous (-) conditions. Gene ontology reveals that the set of genes upregulated in T cells being exposed to porous (-) conditions exhibited enrichment for those encoding proteins involved in diverse molecular functions and cellular processes involving transcription, protein transport, ribosome, cell division, among others (see **Table S1** for a complete list). Moreover, T cells being exposed to porous conditions had higher expression of genes encoding for transcription factors, cytokine receptors and signaling molecules associated with T cell activation. In particular, the TCR signaling pathway was significantly enriched in T cells being exposed to porous surfaces (**Figure 2b** and **Figure S7**).

Among the most significant genes that were upregulated on the porous membrane, CD69 and IL-2 genes stood out as activation markers for T cells. Protein expression (CD69 and IL-2) was measured for both primary human T cells as well as the immortalized human T cell line (Jurkat). Primary human T cells (**Figure S5**) and Jurkat T cells (**Figure 2c**) exhibited a significant increase in expression of the membrane protein CD69 as well as secretion of IL-2 after 24 h of culture on porous compared to non-porous surfaces. Strikingly, up to 20% of the primary human cells exhibited a significant increase in CD69 expression after 24 h, and ~ 10% of the cells expressed CD25 and proliferated after 4 days in conditions promoting formation of microvilli and remarkably, this is happening in the absence of any TCR agonistic triggers (**Figure S5**).

### T cell activation in the absence of TCR agonistic antibodies is independent of surface area

Other studies have shown that in the presence of TCR agonistic antibodies, lymphocyte activation has a higher yield on micropatterned substrates, due to the greater exposure of the cells to a higher density of the activating antibody owing to the larger surface area.^25,26^ To confirm that TCR signaling in the absence of agonistic TCR triggers is induced by nanoconfinement of microvilli, one needs to first assess the possibility of induced stochastic TCR signaling by the greater contact area of the cells with the porous surfaces.

To perform a complete assessment of the relationship between protrusion dimensions and TCR signaling, we used nanoporous membranes with different pore size, interpore distance, pore density and of different materials in absence of TCR agonistic antibodies (**Table 1**). Interestingly, the expression of CD69 and IL-2 secretion by T cells were only enhanced on the porous surface with 200 nm pore size without TCR agonistic antibodies (**Figure 2c**). Moreover, we plotted CD69 expression versus the surface area of the porous membrane, which clearly indicates that there is no correlation between the upregulation of activation markers and the surface area. For example, AAO with 400 nm pores has a 4 times larger surface area compared to AAO with 200 nm pores (200^b^ nm *vs* 400^b^ nm in Table 1), but CD69 and IL-2 were not upregulated on the 400 nm substrates.

**Table 1.** List of the surfaces used for microvilli induction.

| Sample name | material | average pore diameter (nm) | pore density (cm<sup>-2</sup>) | surface area (cm<sup>2</sup>) * |
| --- | --- | --- | --- | --- |
| **20** | AAO | 20 | 10<sup>11</sup> | 84.47853 |
| **100** | AAO | 100 | 10<sup>10</sup> | 42.0686 |
| **200<sup>a</sup>** | Polycarbonate | 200 | 10<sup>7</sup> | 1.40939 |
| **200<sup>b</sup>** | AAO | 200 | 10<sup>9</sup> | 9.2688 |
| **200<sup>c</sup>** | AAO | 200 | 3 x 10<sup>9</sup> | 25.14641 |
| **400<sup>a</sup>** | Polystyrene | 400 | 10<sup>7</sup> | 1.48042 |
| **400<sup>b</sup>** | AAO | 400 | 10<sup>9</sup> | 16.37195 |
| **reference** | Aluminum Oxide or glass | non-porous | 0 | 1.3 |

\* In surface area calculations, it is assumed that 1 $\mu\text{m}$ pore depth is an accessible surface for the microvilli, according to the data in Figure 1e.

Gene transcription analysis revealed that several genes indicating T cell activation – such as *CD69, NFKB1, TCRA, PTPRC* and *ITGAL* – were upregulated on 200 nm pore surfaces. To ask how the pore size tunes the gene expression in T cells, we performed q-PCR analysis on primary human T cells exposed to 200 and 400 nm AAO membranes without TCR agonistic antibodies, 1 hour (early regulation) and 24 hours (late regulation) after seeding (**Figure 2d**). The relative (fold) expression of the target genes was calculated based on the expression of the genes on the non-porous surface as control. qPCR analysis revealed that the target genes were significantly upregulated on 200 nm pores, but not on 400 nm pores. To verify the results, we tested the protein expression of TCR and CD45 at different time points (30 min - 24 hours), which indicated that cells cultured on 200 nm pores exhibited higher expression of TCR and CD45 (**Figure 2d**).

In another attempt, by controlling the depth of the 200 nm pores, we showed that despite an 8-fold increase in the surface area (depth from 0.5 μm to 4 μm), CD69 expression did not change on 200 nm pore membranes (**Figure S5**). These results suggest that 0.5 μm-long protrusion can equally induce enhanced signaling, compared to longer protrusions with an average length of ~ 1.3 μm.

To further establish the correlation between nanoconfined microvilli protrusions and T cell signaling, we perturbed the protrusions with the cytoskeleton-interfering agent Latrunculin B which blocks actin polymerization and observed that CD69 expression was reduced with increasing Lat B concentrations (**Figure S5**). The influence of actin polymerization inhibition on CD69 expression was much more significant on 200 nm pores compared to the other substrates, indicating that the enhanced activation on 200 nm pores is due to microvilli protrusion.

Given the above observations, increased surface area by the pores is not necessarily an improving effect for T cell activation, unless T cells exploit it via microvilli formation. Furthermore, microvilli formation does not necessarily promote T cell signaling, unless its dimensions are in the right range, *i.e*. ~ 200 nm in diameter. We conclude that nanoconfinement of microvilli via infiltration in the porous substrates with 200 nm pore size induces altered cell signaling and potentially T cell activation. But how does imposing a nanoconfinement lead to T cell signaling, particularly in a TCR triggering independent manner?

### Membrane protein sorting by size exclusion

We tested “kinetic segregation” as a model for antibody-independent T cell activation.^27^ In this model, when the inhibitory phosphatase CD45 and TCR are spatiotemporally segregated, a signaling cascade occurs, leading to activation of T cells.^28–31^ When CD45 is absent, sporadic nonspecific interactions lead to temporary TCR chains phosphorylation; in its presence CD45 shields and inhibits nonspecific TCR signals in its proximity. In case of CD45 spatiotemporal exclusion from TCR regions (*e.g*. in “close contacts”^6,7^), sustained downstream TCR signaling and thus T cell activation can take place.^7^ Spatiotemporal segregation is observed to occur both *in vitro* and *in vivo*, even without specific TCR-ligand engagement, however, fast transient triggering does not lead to T cell activation.^7,32,33^

With the above observations, we hypothesized that T cell microvilli nanoconfinement in 200 nm-pores might enforce the spatiotemporal segregation of CD45 and TCR, due to size exclusion. CD45 (~40 nm ^34^) is much larger compared to TCR (~15 nm ^34^), therefore it is plausible that segregation could occur in nanoconfined spaces. In fact, it has been shown in previous studies that physical barriers can hinder diffusion of CD45.^11,24^ More recent studies have shown that in areas of high membrane curvature, such as at the tip of a microvilli, segregation of TCR and CD45 occurs, establishing a phosphatase-free zone, prone for TCR triggering.^5,35^

In our experiments, fluorescence images obtained from Jurkat T cells on porous surfaces demonstrated that the nanoconfinement inhibited or reduced the penetration of large molecules such as CD45 in 200 nm pores (**Figure 3a**). Smaller molecules such as TCR could be found abundantly inside of the microvilli protrusions – recognizable by the dotted structure in the fluorescence images. The induced exclusion of CD45 was not significant in 400 nm pores. The cross-sectional profile of individual microvilli in 200 and 400 nm pores are shown in **Figure S4**. Exclusion of CD45 from 200 nm pores could be caused by several likely synergistic mechanisms. First, the plasma membrane is highly curved around the pore edge and it is well known that high membrane curvature can act as a size-selective diffusion barrier ^36^. Either the high curvature of the plasma membrane around the pore edge creates a size-selective diffusion barrier, as correlations between membrane curvature and proteins are well established ^11,34,37–39^, or the accessible pore volume is largely occupied by the cell membrane and the actin cytoskeleton, leaving little space for the diffusion of proteins with large extracellular domains, such as CD45. Our observations suggest that the confined space of the nanopore enforces spatiotemporal segregation of the signaling complex, and that this phenomenon is completely lost on 1 μm porous membranes (**Figure S4**).

**Figure 3.**
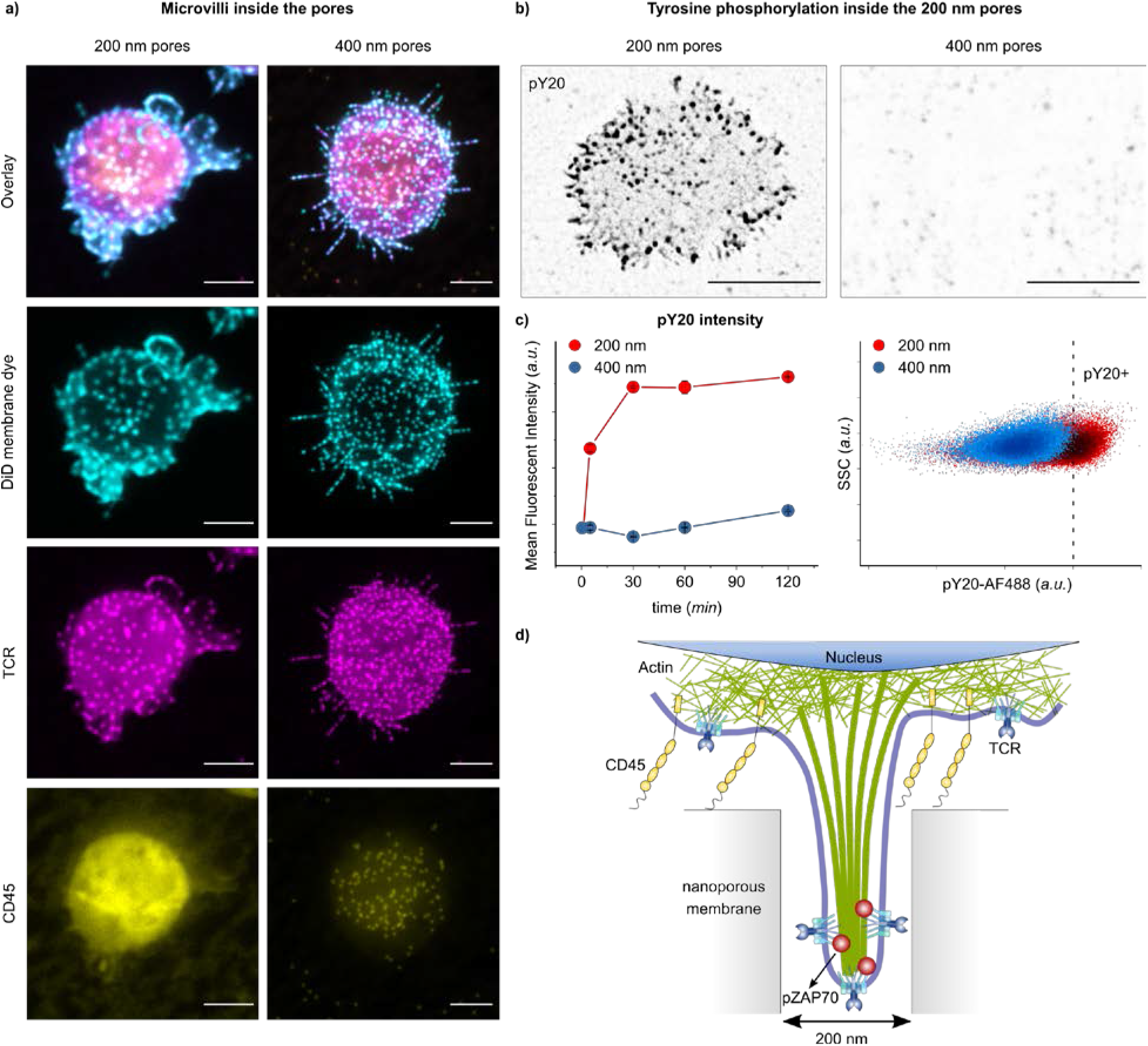
Membrane-bound protein sorting in confined spaces. **a**) shows the projection of confocal fluorescence microscopy images of the cell membrane (stained with DiD), TCR and CD45, from fixed T cells on porous surfaces with (left) 200 nm and (right) 400 nm pore size, 30 min after seeding. Scale bar is 5 μm. **b)** shows the projection of confocal fluorescence microscopy images of pY20, from fixed T cells on 200 and 400 nm pores, fixed 5 min after seeding. The fluorescent signal from inside of the pores is indicated by black dots. Scale bar is 5 μm. **c)** (left) The scatter plot shows the kinetics of the mean fluorescent intensity (MFI) of pY20 of primary human T cells cultured on the indicated surfaces at different time points, measured by flow cytometry. Data represents one of two independent experiments performed in triplicates. Error bars are mean ± s.d. (right) Representative dot plots of pY20 measured by flow cytometry on primary human T cells seeded on 200 nm (red) and 400 nm (blue) porous substrates, 30 min after seeding. **d)** shows schematics of the spatiotemporal segregation model induced by nanoconfinement in 200 nm pores.

To test whether the spatiotemporal segregation is sustained or transient, cells were stained with anti-phosphotyrosine (pY20) as an indicator of TCR signaling (**Figure 3b, c**), as well as pZAP70 (**Figure S6**). Most significantly, fluorescence microscopy revealed the presence of pY signal inside of the 200 nm pores in a dotted pattern, whereas no fluorescent signal for pY was observed inside of the 400 nm pores (**Figure 3b**). In another attempt, the cells were removed from the surface at different time points (5-120 min), and stained with pY20 for flow cytometry analysis (**Figure 3b**). Notably, the cells cultured on 200 nm pores exhibited a strong pY signal. This is rather surprising, because studies on T cell activation on flat surfaces shows that pY signaling is transient and decays within several minutes after activation. The prolonged pY signaling on the porous surface indicates that the nanoconfinement-induced segregation leaves the TCRs accessible to kinases while protecting them from phosphatases that could potentially reverse the phosphorylation, allowing for sustained signaling during their residence inside of the pores (**Figure 3d**).

### Augmented and sustained ERK phosphorylation induced by nanoconfinement

Next we asked whether TCR phosphorylation inside of the 200 nm pores can initiate downstream signaling to activate T cells with/without TCR agonistic antibodies. Therefore, we looked into the extracellular-signal-regulated kinase (ERK) pathway in T cells exposed to 200 nm porous surfaces and compared it with agonistic TCR antibody-activated T cells as a reference. The ERK pathway is part of the mitogen-activated protein kinase (MAP Kinase) pathway, a major signaling cascade in regulation of T cell activation, differentiation and proliferation^40^. ERK signaling is activated during antigen recognition by T cells, where the strength of the signal and its duration depend on the affinity and avidity of the TCR-antigen interactions.^32,41^ Additionally, recent studies propose a correlation between ERK phosphorylation and podosome maturation (*i.e*. nanoscale actin-rich protrusions).^42–44^

The cells were cultured on either TCR agonistic antibody (*i.e*. αCD3/αCD28) coated (+) and non-coated (-) porous (200 nm) or non-porous surfaces. T cells activated on AAO with 200 nm pores with TCR agonistic antibody (shown as porous (+)) exhibited a significantly higher phosphorylation of ERK, which was sustained much longer than in cells activated on the non-porous (+) surface (**Figure 4a, b**). ERK signaling in non-porous (+) cells peaked at 15 min and ceased in most cells after 60 min. On the other hand, porous (+) cells had a higher amount of ERK phosphorylation (pERK) per cell (as measured by mean fluorescent intensity, MFI), which was sustained for longer than in non-porous (+) cells. Interestingly, cells cultured on AAO with 200 nm pores with TCR agonistic antibody (shown as porous (-)) also exhibited amounts of pERK comparable to those in non-porous (+) cells. ERK was not activated in resting cells on the non-porous (-) surfaces (**Figure 4a**, the control experiment).

**Figure 4.**
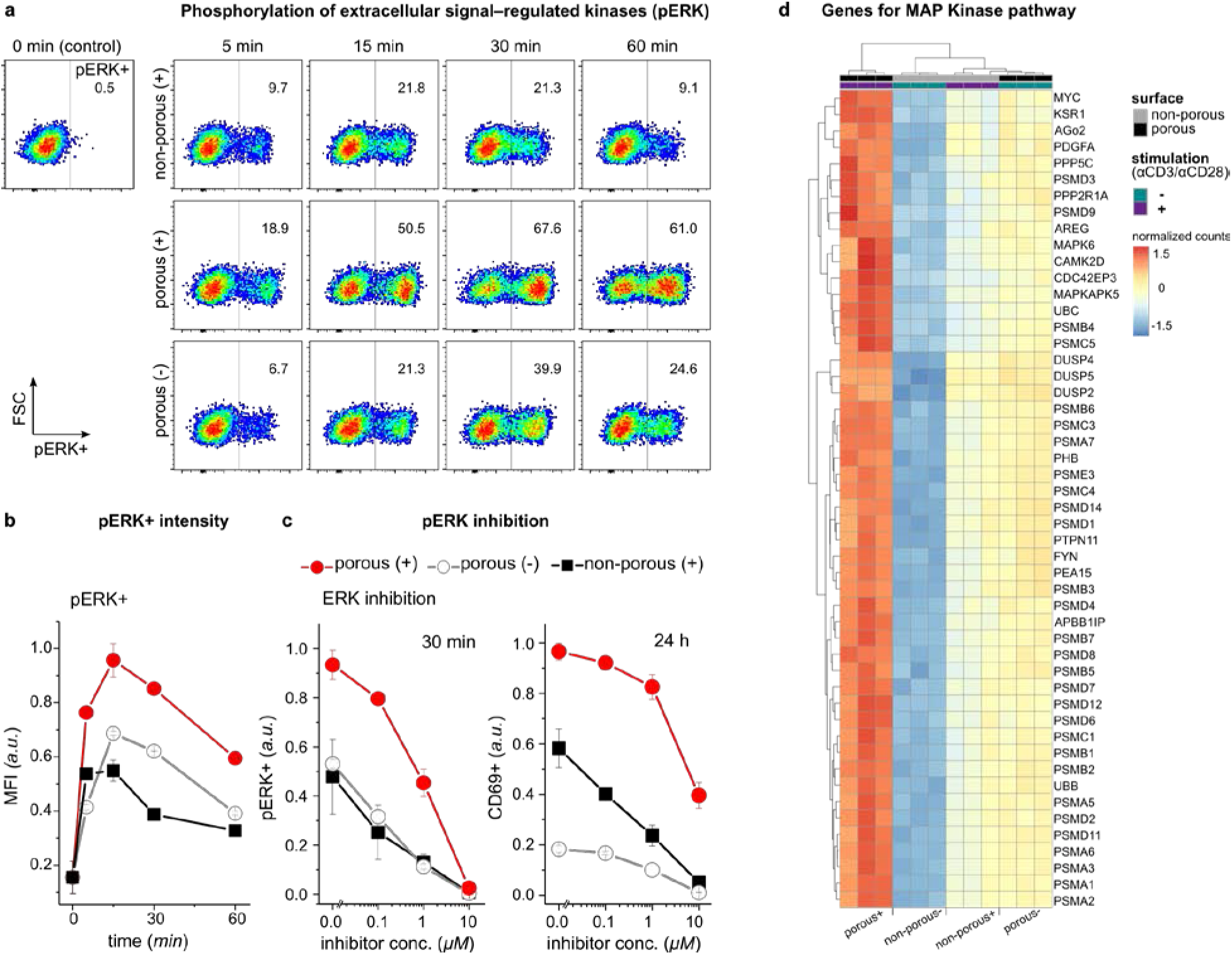
ERK phosphorylation is augmented and sustained on the porous membranes. **a)** Representative dot plots of phosphorylated ERK measured by flow cytometry on Jurkat T cells under the indicated conditions. Data is representative from two independent experiments performed in duplicates or triplicates. **b)** The bar diagram shows the kinetics of the mean fluorescent intensity (MFI) of ERK-phosphorylated cells (pERK+) with the indicated treatment. Data represents one of two independent experiments performed in duplicates or triplicates. Error bars are mean ± s.d. **c)** Jurkat T cells were incubated with ERK inhibitor U0126 in different concentrations for 30 min before being activated on the indicated surfaces. Bar diagrams show percentages of pERK+ cells after 30 min and CD69+ cells after 24 hours measured by flow cytometry (n = 3 replicates). Error bars are mean ± s.d. **d)** Heatmap of the top 50 significantly upregulated genes in the MAP Kinase pathway of primary human T cells activated on the indicated surfaces for 4 h, adjusted P values < 0.05. Genes were selected based on the comparison between porous (+) and non-porous (-) cells (n = 3 replicates). The complete set of the significantly differentially expressed genes in MAP Kinase pathway can be found in the **supplementary information**.

To further elucidate the correlation between nanoconfined microvilli protrusions and ERK signaling, we used ERK signaling inhibitor U0126 before culturing the cells on the surfaces. As expected, we observed reduced ERK phosphorylation in a concentration-dependent fashion (**Figure 4c**). ERK inhibition correlated with reduced percentages of CD69+ cells 24 h after activation in all conditions. Moreover, pathway enrichment analysis for differentially expressed genes indicated that the MAP Kinase pathway was enriched in all of the three groups (*i.e*. porous (+), porous (-) and non-porous (+)) compared to non-porous (-), as shown in **Figure 4d** and **Figure S8**). There was a large overlap in the corresponding gene expression patterns between porous (-) and non-porous (+) cells, whereas expression levels were much higher on porous (+) cells. These results suggest that nanotopographical constraints have a central role in the enhanced and sustained MAP Kinase signaling.

### T cell activation is boosted by nanotopographical constraints

The process of T cell activation and signaling in response to the physical constraints can be exploited for many applications, including the topographic design of biomaterials for enhanced activation and proliferation of the cells.^45–49^ We examined the suitability of our 200 nm pore platform for Pan T cell activation, to validate its potential for application in immunotherapy, by enhancing activation of T cells via imposing a physical constraint (**Figure 5**). For polyclonal T cell activation and expansion, the surfaces of porous and non-porous aluminum oxide films were coated using activating antibodies against CD3 (αCD3; TCR stimulus) and CD28 (αCD28; costimulatory cue).

**Figure 5.**
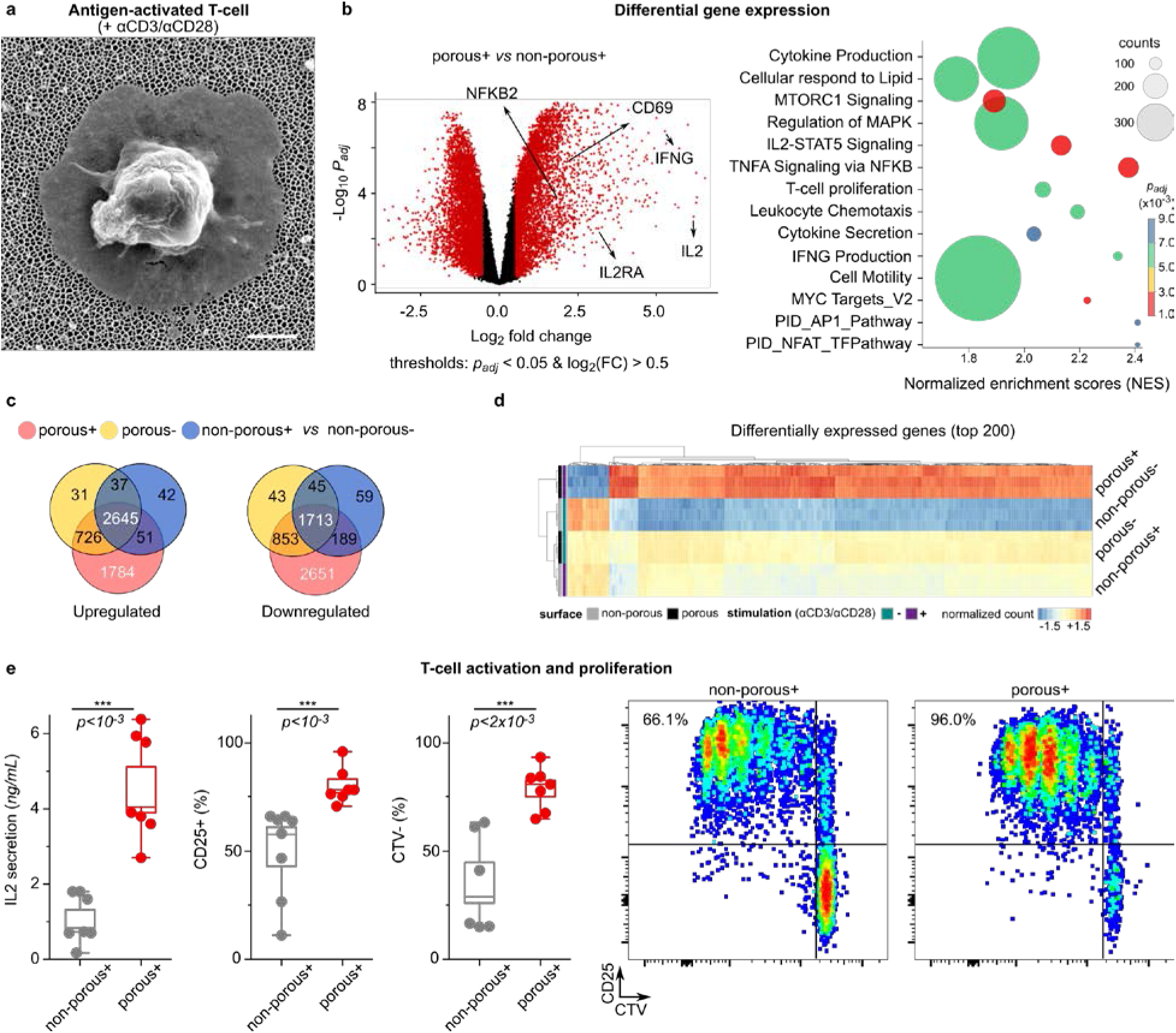
Boosted T cell activation and proliferation by combining nano-topographical and biochemical cues. **a)** Scanning electron microscope (SEM) image of an activated primary human T cell on top of the nanoporous membrane with 200 nm pore size. Scale bar 1 μm. **b)** (left) Volcano plot showing differences in RNA expression of T cells activated with antibodies on porous (+) vs non-porous (+) surfaces. Red color indicates significance with adjusted *P* value <0.05, and log2(fold changes) > 0.5. (right) Pathway enrichment of differentially expressed genes in porous (+) cells versus non-porous (+) T cells shows increased stimulation of porous (+) T cells. The size of the circle corresponds to the gene counts (from the reference pathway), the color corresponds to the adjusted *P* value. **c)** Venn diagrams of the differentially expressed genes (upregulated and downregulated) in three conditions (porous (+), porous (-) and non-porous (+) compared to the non-porous (-) cells). **d)** The top 200 significantly upregulated genes in porous (+) compared to the non-porous (-) cells, based on adjusted *P* values < 0.05 (n = 3 replicates). +/- signs indicate the presence of activation antibodies (αCD3/CD28) on the surface. The complete set of the significantly differentially expressed genes can be found in the **supplementary information**. **e)** Box diagrams show activation of human primary T cells on porous surfaces. IL-2 secretion was measured after 24 hours, and CD25 expression was measured after 4 days. Three independent experiments were performed in duplicates or triplicates. The p-values were determined by two-sided Mann-Whitney tests in R. The proliferation assay (CTV) was used to assess the expansion of the cells after 4 days. The graphs represent examples of the measurements by flow cytometry.

Differential gene expression analysis revealed that more than 2700 genes were upregulated in T cells cultured on porous (+) (*i.e*. AAO 200 nm now in the presence of TCR agonistic antibodies), compared to cells cultured on non-porous (+) surfaces, whereas more than 2300 genes were downregulated. Venn diagrams of the differentially expressed genes (compared to non-stimulated cells) indicate the significance of the nanotopography-induced gene regulation. The list of the differentially expressed genes is provided in supplementary information (**Table S1**). Gene ontology comparison showed that the set of genes upregulated by porous (+) compared to non-porous (+) surfaces, are enriched for biological processes such as regulation of cellular amino acid metabolic process, NIK/NF-kappaB signaling, T cell receptor signaling pathway and cell division (**Figure S7**).

After activation with TCR agonistic antibodies, T cells exhibited a significant increase in expression and secretion of IL-2 after 24 h of culture on porous compared to non-porous surfaces (**Figure 5e**). Higher IL-2 levels correlated with more proliferating primary T cells with high CD25 expression, as measured 4 days after activation via a dye dilution assay (CTV) (**Figure 5e**). The observed boosted T cell response on 200 nm pores is therefore due to the concurring biochemical and biomechanical signals that are induced by TCR activation and nanotopographical constraints, respectively. Moreover, it is worth pointing out that cells activated on porous and non-porous share similar phenotypic characteristics despite enhanced activation and proliferation (**Figure S9**).

## Conclusions

In summary, we show here for the first time that nanoporous membranes with 200 nm pores promote the creation of TCR nanoclusters at the tips of the microvilli protrusions, thereby boosting T cell signaling, activation and proliferation. TCR-independent T cell signaling is facilitated via pore size-dependent segregation of membrane proteins, amplifying and sustaining the necessary signaling events that lead to T cell activation in absence of cognate antigen-mediated T cell activation. We discovered that kinetic segregation can be locally induced when T cells protrude into nanoporous structures. Notably, cell signaling cascades and gene regulation are maximally altered in cells cultured on nanoporous membranes with 200 nm pores, including ERK phosphorylation as a component of MAP Kinase pathway and T cell activation. Future studies will need to address what other proteins/molecules might be involved in the membrane-curvature-induced segregation of the TCR and CD45 phosphatase signaling complex and to what extent they resemble the previously described microsynapses/podosynapses that are enriched in TCR signaling molecules.^1,50^ Our discovery is of fundamental importance in immunology and cell biology, because we established a direct causality between size-dependent microvilli formation and markedly altered gene expression in T cells, which supports T cell activation. Also the synergistic combination of protrusion formation and antibody stimulation presents a simple strategy to optimize *in vitro* T cell activation and may be of value for protocols for adoptive immune cell therapy.

## Data availability

All the data needed to evaluate the conclusions in the paper are present in the paper. Additional data and other findings of this study are available from the corresponding authors upon reasonable request.

## Acknowledgments

This work was supported by the Human Frontiers Science Program RGY0065/2017 to E.K. This project received funding from Swiss National Science Foundation Spark grant CRSK-3_190394 to M.A. and D.S. M.A. is thankful to The Holcim Foundation for the Promotion of Scientific Training, for partially funding the project.

We are grateful to members of our laboratories for assistance during the project, in particular Lina Aires, Johanna Mehl, Arianna Menghini, Cameron Moshfegh, Lion Raaz, Norma De Giuseppe, Chantel Spencer-Hänsch and Isabel Gerber. The authors acknowledge the valuable and insightful discussions with János Vörös, Jonas Ries, Michael Dustin and Gerhard Schütz. We thank Daniel Hoces and Leonhard Sigel for the valuable clinical assistance. We are thankful to Tobias Wolf and Federica Sallusto for the discussions as well as materials. The authors acknowledge support of the Scientific Center for Optical and Electron Microscopy ScopeM of the Swiss Federal Institute of Technology ETHZ and Tobias Schwarz. We thank the FGCZ (Functional Genomics Center Zurich) at ETH Zurich and the University of Zurich (UZH), as well as Emilio Yángüez and Ge Tan for conducting the high throughput RNA-seq.

## Contributions

M.A., D.S. and E.K. conceived the project and designed the experiments. M.A., D.S., T.Z., J-Y.S, M.AS., M.B. and S.L. performed the experiments. I.S. and M.A. carried out the bioinformatics data analysis of RNA-seq. S.J.I, C.F. and M.A. conducted the image analyses. D.S., A.O., V.V. and E.K. contributed to concept advancement, data analysis and manuscript preparation. All authors contributed in analysis and interpretation of data. M.A. wrote the manuscript with support from all authors.

## Ethics declarations

The authors declare no competing interests.

## Notes

### Competing Interest Statement

The authors have declared no competing interest.

